# Machine learning models for prediction of xenobiotic chemicals with high propensity to transfer into human milk

**DOI:** 10.1101/2023.08.06.552173

**Authors:** Sudharsan Vijayaraghavan, Akshaya Lakshminarayanan, Naman Bhargava, Janani Ravichandran, R.P. Vivek-Ananth, Areejit Samal

**Author notes:** Corresponding authors (R.P. Vivek-Ananth); (Areejit Samal).

## Abstract

Breast milk serves as a vital source of essential nutrients for infants. However, human milk contamination via transfer of environmental chemicals from maternal exposome is a significant concern for infant health. Machine learning based predictive toxicology models can be valuable in predicting chemicals with high propensity to transfer into human milk. To this end, we build such classification- and regression-based models by employing multiple machine learning algorithms and leveraging the largest curated dataset to date of 375 chemicals with known Milk to Plasma concentration (M/P) ratios. Our Support Vector Machine (SVM) based classifier outperforms other models in terms of different performance metrics, when evaluated on both (internal) test data and external test dataset. Specifically, the SVM based classifier on (internal) test data achieved a classification accuracy of 77.33%, specificity of 84%, sensitivity of 64%, and F-score of 65.31%. When evaluated on an external test dataset, our SVM based classifier is found to be generalizable with sensitivity of 77.78%. While we were able to build highly predictive classification models, our best regression models for predicting the M/P ratio of chemicals could achieve only moderate R^2^ values on the (internal) test data. As noted in earlier literature, our study also highlights the challenges in developing accurate regression models for predicting the M/P ratio of xenobiotic chemicals. We have made our complete workflow, train and test datasets, and computer codes for the classification and regression models publicly available via a dedicated GitHub repository. Overall, this study attests the immense potential of predictive computational toxicology models in characterizing the myriad chemicals in the human exposome.

## 1. Introduction

Breast milk is widely recognized as the optimal source of nutrition for infants and provides numerous benefits to both the infant and the mother. Published studies, including by Rollins *et al*. (Rollins et al., 2016), have shown that breastfeeding contributes towards a world that is healthier, better educated, more equitable and more environmentally sustainable. Breast milk is also known to provide protection to the infant from health complications such as cardiovascular disease, sudden infant death syndrome, growth faltering, and inflammatory bowel disease (Hauck et al., 2011). Notably, breastfeeding has been shown to be associated with reduced risk in mothers for premenopausal breast cancer, ovarian cancer, retained gestational weight gain, type 2 diabetes, myocardial infarction, and metabolic syndrome (Stuebe, 2009). Moreover, breast milk is an eco-friendly and cost-effective option compared to using infant formula (Bartick et al., 2017).

In spite of the benefits associated with breastfeeding, there is a legitimate concern about the potential exposure of infants to environmental chemicals (including drugs) through lactation (Karthikeyan et al., 2021; Lehmann et al., 2018). Pregnant women and lactating mothers are exposed to a wide range of environmental chemicals via food, medication, personal care products, and environmental pollutants (Landrigan et al., 2002; Mead, 2008; Sonawane, 1995). Exposure to such chemicals can affect the health of both the mother (Li et al., 2020) and the development of breastfed infants (Leibson et al., 2018). Wild (Wild, 2005) proposed the concept of ‘exposome’ to describe the non-genetic factors that influence health and disease of an individual, starting from the prenatal period. In essence, exposome captures the sum total of all environmental exposures that an individual experiences through their life, and the associated health effects (Miller and Jones, 2014; Rappaport and Smith, 2010; Vermeulen et al., 2020; Wild, 2005). Consequently, due to potential impact on infant and maternal health, there is significant interest in characterizing environmental chemicals with high propensity to transfer from maternal plasma to human milk, i.e. potential human milk contaminants in the chemical exposome (Karthikeyan et al., 2021; Lehmann et al., 2018). In this direction, some of the authors of this study had previously built an online knowledgebase, ExHuMId (Karthikeyan et al., 2021), that compiles experimentally detected human milk contaminants from published studies analyzing breast milk samples from India.

Notably, experimentally measured Milk to Plasma concentration (M/P) ratio is used to identify the equilibrium concentration of a chemical in maternal plasma in comparison to breast milk, and this M/P ratio is used as an indicator for the propensity of a xenobiotic chemical to enter human milk (Agatonovic-Kustrin et al., 2002; Anadón et al., 2011; Vasios et al., 2016). In terms of environmental chemicals in the human exposome, several studies have shown lipophilic substances to have a high potential to transfer into human milk from maternal plasma via passive diffusion (Agatonovic-Kustrin et al., 2002; Anadón et al., 2011; Heinzow, 2009; Vasios et al., 2016; Zhao et al., 2006). Further, a systematic analysis (Karthikeyan et al., 2021) based on available M/P ratios for human milk contaminants in ExHuMId revealed structural properties of chemicals that can influence their transfer into human milk. In a nutshell, previous analyses have led to realization that physicochemical properties of xenobiotic chemicals influence their potential to transfer from maternal plasma to human milk (Agatonovic-Kustrin et al., 2002; Anadón et al., 2011; Heinzow, 2009; Karthikeyan et al., 2021; Vasios et al., 2016; Zhao et al., 2006).

Experimental measurement of a xenobiotic chemical’s propensity to enter human milk is both a difficult task and ethically impractical. Since the 1980s (Begg and Atkinson, 1993; Wilson et al., 1980), there have been attempts to predict M/P ratio for xenobiotic chemicals, and the initial studies were based on methods incorporating physicochemical properties of the chemicals while ignoring clinical information or effects of active transport. Following the initial attempts, several studies employing machine learning algorithms and quantitative structure-activity relationship (QSAR) principle have been proposed (Abraham et al., 2009; Agatonovic-Kustrin et al., 2000; Fatemi and Ghorbanzad’e, 2010; Kar and Roy, 2013; Katritzky et al., 2005; Wanat et al., 2020; Yap and Chen, 2005). Such predictive models have been built on the QSAR principle that the biological or chemical activity of a compound can be quantitatively related to its molecular structure and physicochemical properties. For instance, Agatonovic-Kustrin *et al*. (Agatonovic-Kustrin et al., 2000) developed a genetic neural network based model to predict the degree of drug transfer into breast milk using a dataset of 60 drugs and their experimentally derived M/P ratios, and the authors reported a coefficient of determination (R^2^) value > 0.96 on train data and a root mean square (RMS) value of 0.425 on test data for their best model. Yap and Chen (Yap and Chen, 2005) developed a regression model based on general regression neural network by using a dataset of 102 chemicals in train data and 20 chemicals in test data, and the authors reported a R^2^ value of 0.677 and mean squared error (MSE) value of 0.206 on test data for their best model. Katritzky *et al*. (Katritzky et al., 2005) built a predictive model by dividing a dataset of 100 chemicals into three subsets. Thereafter, the authors created three training datasets by combining any two subsets, followed by derivation of equations for each training dataset. The authors then employed these equations to predict the logarithm of M/P ratios for the chemicals in the corresponding test data. For the developed models, Katritzky *et al*. (Katritzky et al., 2005) reported an average R^2^ value of 0.763. Abraham *et al*. (Abraham et al., 2009) employed an artificial neural network to predict the logarithmically transformed M/P ratios of chemicals by using a dataset of 179 drugs and environmental pollutants of which 135 chemicals were in the train data, 22 chemicals in the test data, and further 22 chemicals in the external test data. For the best model, Abraham *et al*. (Abraham et al., 2009) reported root mean square error (RMSE) of 0.056 on train data, 0.109 on test data, and 0.09 on external test data. However, Abraham *et al*. (Abraham et al., 2009) did not report the R^2^ values for their model. Fatemi and Ghorbanzad’e (Fatemi and Ghorbanzad’e, 2010) developed a counter propagation artificial neural network based classification model using a dataset of 124 chemicals in the train data, 20 chemicals in the test data, and 10 chemicals in the external test data, and the authors reported an accuracy of 100% on train data, 100% on test data, and 90% on external test data. Kar and Roy (Kar and Roy, 2013) developed both classification and regression models using linear discriminant analysis (LDA) and multiple linear regression (MLR), respectively, using a dataset consisting of 97 chemicals in the train data and 88 chemicals in the test data. For their regression model, the authors obtained a R^2^ value of 0.7 on train data, and for their classification model, the authors obtained an accuracy of 88.66% and 56.82% on train and test data, respectively, sensitivity of 82.05% and 63.16% on train and test data, respectively, and F-score of 85.33% and 55.81% on train and test data, respectively. However, Kar and Roy (Kar and Roy, 2013) did not report the R^2^ value for their regression model on test data. Wanat *et al*. (Wanat et al., 2020) developed three regression models to predict the M/P ratio of chemicals, namely, MLR, partial least squares (PLS), and random forest (RF). This study identified MLR and RF as the most effective models with MLR achieving R^2^ value of 0.86 on a train data of 28 chemicals while RF achieving R^2^ value of 0.86 on a train data of 58 chemicals and 0.29 on test data of 25 chemicals.

Despite the availability of accurate models, the prediction of transfer into milk from maternal plasma for actively transported drugs remains a challenging endeavour (Anderson and Momper, 2020). Further, the applicability and generalizability of the previously published machine learning models for this purpose is hindered by the limited sizes of the train and test datasets used to build them. Moreover, a minority of the previously published models have made their train and test dataset of chemicals accessible in the associated publication, and none of the previously published models have made the computer codes available. Thus, it is very difficult to reproduce the results from the previously published models on this topic.

The primary aim of this study is to build accurate machine learning models for predicting high propensity of transfer of xenobiotic chemicals from maternal plasma to human breast milk by leveraging, to date, the largest curated dataset of chemicals with experimentally determined M/P ratios. Notably, our approach upholds the three essential principles of data science: repeatability, reproducibility, and replicability. To accomplish this, we manually curated a dataset of 375 chemicals along with their experimentally determined M/P ratios from previously published scientific literature (Supplementary Table S1). Thereafter, we computed 1875 molecular descriptors for each chemical in our dataset, and the computed descriptors capture the structural and physicochemical features of the chemical dataset. Subsequently, we utilised the computed descriptors and known M/P ratio for chemicals as features along with multiple machine learning algorithms to build predictive models for the purpose. Our workflow (Figure 1) led to reliable classification models for predicting chemicals with ‘high risk’ of transfer from maternal plasma to human milk. Although we encountered challenges in building accurate regression based models to predict the M/P ratio of chemicals, we were able to partially address these challenges by drawing insights from previous work and were able to build a regression model that gave reasonable and better results in comparison to models proposed earlier. Interestingly, our regression models yielded favourable results when challenged with the classification task. Lastly, to assess the applicability and generalizability of the built models, we validated our classification models by leveraging a large external test dataset of 202 chemicals (Supplementary Table S2), and this evaluation on an external test dataset highlighted the robustness of the results obtained in this study. Overall, this study also takes a step towards achieving FAIR (Wilkinson et al., 2016) compliance by publicly releasing our curated chemical datasets and computer codes through the associated GitHub repository (https://github.com/asamallab/M-by-P-ratio-Pred), and this is expected to enable wider scientific community to reproduce, use and build upon our work. By emphasizing on repeatability, reproducibility and replicability in this research, we hope to contribute to the recent thrust in the field of computational toxicology towards creating transparent and dependable models.

**Figure 1:**
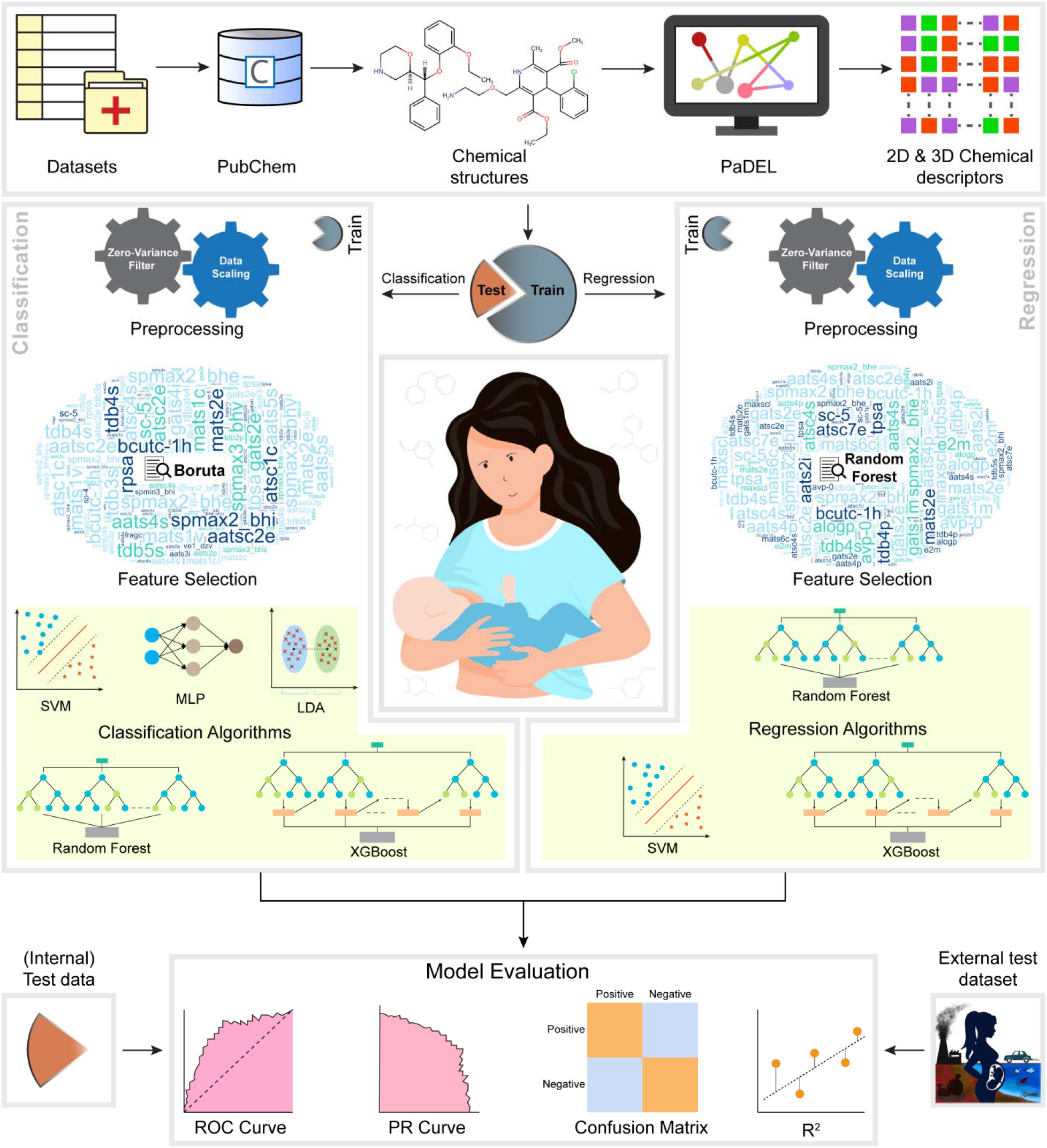
Schematic diagram summarizing the workflow to build the classification- and regression-based machine learning models to predict xenobiotic chemicals with high propensity to transfer from maternal plasma to human milk. This figure shows the key steps involved in data curation, feature generation, data preprocessing, feature selection, and the training and evaluation of classification- and regression-based machine learning models. The word clouds in this figure were generated using https://www.wordclouds.com.

## 2. Methods

### 2.1 Chemical dataset

In this study, we compiled from published literature a curated dataset of 375 chemicals (Supplementary Table S1), consisting of xenobiotic chemicals (mainly, drugs), with known experimentally determined Milk / Plasma concentration (M/P) ratios. Majority (368) of the chemicals in our dataset were compiled from Vasios *et al*. (Vasios et al., 2016) and information on few additional chemicals were obtained from other publications (Fatemi and Ghorbanzad’e, 2010; Ito et al., 2015; Larsen et al., 2003; Wanat et al., 2020). For 4 chemicals in our dataset, the M/P ratios were available from multiple publications, and in such cases, the mean of the reported M/P ratios for a chemical across publications was used. In a few instances, we did not include chemicals with known M/P ratio in the published literature as the computation of the molecular descriptors (or features) failed for them.

For our curated dataset of 375 chemicals with known M/P ratios, we obtained the two-dimensional (2D) structures from PubChem (Kim et al., 2023). We converted the 2D structures of these 375 chemicals to their corresponding three-dimensional (3D) structures as follows. Using RDKit (Landrum, 2022), we first embed a chemical in 3D space by employing the ETKDG method. Thereafter, the 3D structure of a chemical was energy minimized using MMFF94 forcefield in RDKit. Subsequently, we computed the molecular descriptors for the 375 chemicals in our dataset using PaDEL version 2.21 (Yap, 2011). A total of 1875 molecular descriptors were computed for each chemical using PaDEL including topological, geometric, electrostatic, and several other types of descriptors. Note that the 2D structure of a chemical was used to compute one-dimensional (1D) and 2D descriptors, while the 3D structure was used to compute 3D descriptors in PaDEL. The computed descriptors were used as features to build machine learning models.

In this study, we build both classification and regression models to predict the propensity of a chemical to transfer from plasma to milk. For the classification models, chemicals with M/P ratio ≥ 1 were designated as ‘high risk’ while the remaining chemicals with M/P ratio < 1 were designated as ‘low risk’, following previously published studies (Fatemi and Ghorbanzad’e, 2010; Kar and Roy, 2013; Vasios et al., 2016). For the regression models, we treated the M/P ratio of chemicals in our dataset as a continuous variable. A logarithmic transformation was used to reduce the skewness in the distribution of M/P ratios for chemicals in our dataset. Note that a constant value of 1 was added to the M/P ratio of each chemical in our dataset before logarithmic transformation in order to avoid undefined logarithmic values for chemicals with M/P ratio equal to 0.

To build the classification and regression models, our dataset of 375 chemicals (250 chemicals categorized as ‘low risk’ and 125 chemicals categorized as ‘high risk’) with experimentally determined M/P ratios was randomly split into two sets with 80% of the data in the train set and the remaining 20% of the data in the test set (Supplementary Table S1). Note that the initial dataset of 375 chemicals exhibited an imbalanced distribution of chemicals, with a 2:1 ratio between ‘low risk’ and ‘high risk’ classes. During the process of splitting the data into train and test set, we preserved the ratio between the ‘low risk’ and ‘high risk’ chemicals in both train and test set.

In addition to this train and test dataset (together consisting of 375 chemicals with M/P ratios), we also compiled an external test dataset of 202 chemicals from Karthikeyan *et al*. (Karthikeyan et al., 2021), Lehmann *et al*. (Lehmann et al., 2018) and Neveu *et al*. (Neveu et al., 2017), which have been experimentally detected in human breast milk samples (Supplementary Table S2). Since the 202 chemicals in the external test dataset have been experimentally detected in human breast milk samples, they were categorized as ‘high risk’ chemicals, and thereafter, used for the validation of classification models. However, the M/P ratio for these 202 chemicals in the external test dataset have not been reported in the literature, and thus, the external test dataset cannot be used to evaluate the regression models. For the 202 chemicals in the external test dataset, we obtained the 2D structures from Karthikeyan *et al*. (Karthikeyan et al., 2021). Thereafter, we followed the same procedure as described above for the 375 chemicals to generate the 3D structures and compute the 1875 molecular descriptors for the 202 chemicals in the external test dataset.

### 2.2 Feature selection

Each chemical in our dataset has 1876 features including the dependent variable (M/P ratio). After randomly splitting the dataset of 375 chemicals into the train set with 80% of the data and the test set with the remaining 20% of the data, we first removed the features with zero variance in the train set. Thereafter, the train and test sets were scaled using Standardscaler in scikit-learn (Pedregosa et al., 2011).

While building classification models, we used BorutaPy, a wrapper built on a random forest-based classifier, to implement feature selection (Daniel, 2022; Kursa and Rudnicki, 2010). Notably, BorutaPy takes into account multi-variable relationships and considers all features that are relevant for the dependent variable. The method involves taking a copy of the original features and shuffling them to create shadow features. These shadow features are concatenated with the train set to find the importance of individual features using Z-score (Kursa and Rudnicki, 2010). If the importance of a feature is greater than the maximum importance of the shadow features, then such a feature is retained. Otherwise, the feature is considered unimportant and dropped while building the classification models. While building regression models, we used the random forest method (Breiman, 2001) within scikit-learn to select the important features.

### 2.3 Machine learning algorithms

In this study, we implemented several machine learning algorithms namely, support vector machine (SVM), extreme gradient boosting (XGBoost), linear discriminant analysis (LDA), multi-layer perceptron (MLP), and random forest (RF), to build classification and regression models. In this subsection, we provide a concise overview of the algorithms employed here.

#### 2.3.1 Support vector machine

Support vector machine (SVM) is a supervised machine learning algorithm used for classification and regression tasks. It maximizes predictive accuracy and addresses overfitting by finding an optimal hyperplane that maximizes the margin between training examples and class boundaries (Boser et al., 1992; Cortes and Vapnik, 1995; Drucker et al., 1996). SVM can handle both binary and multi-class classification tasks, and can also handle non-linear data by introducing a new dimension. The robustness and accuracy of such models can be improved through hyperparameter tuning. Additionally, there is a trade-off between correctly classified points and maximizing the margin. Support vector regressor (SVR) is a similar technique used for regression tasks, and it aims to find the hyperplane that maximizes the margin distance between data points by approximating the relationship between inputs and continuous targets. SVR is particularly effective for handling non-linear and high-dimensional data.

#### 2.3.2 Extreme gradient boosting

Extreme gradient boosting (XGBoost) is a widely-used machine learning algorithm for classification and regression tasks (Friedman, 2001). It combines weak learners, known as base learners, to create a stronger learner by iteratively minimizing the overall error. XGBoost offers advantages such as high accuracy, fast execution speed, regularization techniques, and flexibility compared to other popular algorithms (Chen and Guestrin, 2016).

#### 2.3.3 Linear discriminant analysis

Linear discriminant analysis (LDA) is a computationally efficient machine learning algorithm used for classification. LDA optimizes class separation by maximizing variance between classes and minimizing variance within classes through a linear discriminant function. It assumes Gaussian probability density functions with the same covariance for each class. LDA projects data onto a lower-dimensional space by maximizing the distances between the means of the classes while minimizing the within-class variance (Fisher, 1936). LDA has advantages such as handling high-dimensional data, generative modeling, and dimensionality reduction. However, the method has limitations such as assumptions of linearity and normal distribution of data, which may not hold in all cases (Tharwat et al., 2017).

#### 2.3.4 Multi-layer perceptron

Multi-layer perceptron (MLP) is a neural network used for classification and regression tasks. It consists of multiple layers of interconnected nodes or neurons, forming a feedforward network. Typically, a MLP has input, hidden and output layers (Hinton, 1990). Each node in a layer is connected to every node in the subsequent layer, with associated weights that are updated during training. For classification, the MLP predicts the probability of input belonging to each class, with the highest probability determining the final output. MLPs are popular for their ability to learn complex non-linear relationships between input features and output variables (Rumelhart et al., 1986). However, this algorithm can be computationally expensive when modeling large datasets.

#### 2.3.5 Random forest

Random forest (RF) is a versatile machine learning algorithm used for classification and regression tasks. It consists of multiple tree predictors, where each tree’s outcome is influenced by a random vector sampled independently across all the trees in the forest (Breiman, 2001). In classification, the class predicted by the majority of trees determines the final result, while in regression, the final prediction is the average of predictions from all the trees. RF excels in handling high-dimensional datasets and produces stable predictions (Breiman, 2001; Liaw et al., 2002). However, the computational cost of this algorithm increases when modeling large datasets.

### 2.4 Hyperparameter tuning in algorithms

Hyperparameter tuning is a technique used to exhaustively search for the best parameters among a given set of parameters for an algorithm to build predictive models. Hyperparameters of an estimator (algorithm) are a set of parameters that control the learning of the train data and determine the performance of the model on the test data. A notable challenge during model selection is to find the right combination of the hyperparameters for an estimator from the available hyperparameter space. In this study, we used GridSearchCV in scikit-learn (Pedregosa et al., 2011) for hyperparameter tuning. In GridSearchCV, for an estimator and a set of hyperparameters, we performed hyperparameter tuning using repeated 10-fold cross-validation. Subsequent to hyperparameter tuning, the final model is trained based on the best set of parameters obtained from GridSearchCV. Supplementary Table S3 lists the best sets of parameters obtained from GridSearchCV for different estimators used in this study. Supplementary Table S4 lists the final set of top ranked features which are used to build the classification and regression models in this study.

### 2.5 Domain of Applicability

In Setubal workshop report (Jaworska et al., 2005), the applicability domain (AD) of a QSAR model is defined as: “The AD of a (Q)SAR is the physico-chemical, structural, or biological space, knowledge or information on which the training set of the model has been developed, and for which it is applicable to make predictions for new compounds. The AD of a (Q)SAR should be described in terms of the most relevant parameters, i.e. usually those that are descriptors of the model. Ideally, the (Q)SAR should only be used to make predictions within that domain by interpolation not extrapolation”. In other words, the prediction made by a QSAR model is reliable or acceptable only if the chemical in the test data lies in the applicability domain of the model. Therefore, the domain of applicability can be used to better understand the scope of a predictive model.

In this study, we employed the standardized approach proposed by Roy *et al*. to find whether the chemicals in our test data fall within the applicability domain of the built models for classification and regression (Roy et al., 2015).

## 3. Results

### 3.1 Workflow for building classification and regression models

The aim of this study is to build machine learning models to predict xenobiotic chemicals with high propensity to transfer from maternal plasma to human milk. Firstly, we build classification-based models that can accurately categorize chemicals as either ‘high risk’ or ‘low risk’ of transfer from maternal plasma to human milk. Secondly, we build regression-based models that can predict the Milk / Plasma concentration (M/P) ratios for xenobiotic chemicals. Figure 1 presents a schematic workflow of the different steps undertaken in this study to build the classification- and regression-based machine learning models.

To build the machine learning models, we leveraged a curated dataset of 375 chemicals with experimentally determined M/P ratios compiled from Vasios *et al*. (Vasios et al., 2016) and other published literature (Fatemi and Ghorbanzad’e, 2010; Ito et al., 2015; Larsen et al., 2003; Wanat et al., 2020) (Methods; Supplementary Table S1). For each chemical in this dataset, we obtained the 2D structure, generated the 3D structure, and computed 1875 molecular descriptors using PaDEL (Yap, 2011). The 1875 molecular descriptors for a chemical are used as features in the machine learning models (Methods). The curated dataset of 375 chemicals with M/P ratios was randomly divided into two sets namely, a train set with 80% of the data and a test set with the remaining 20% of the data, for building the predictive models (Methods).

To build the classification models, we employed five different machine learning algorithms namely, support vector machine (SVM), extreme gradient boosting (XGBoost), linear discriminant analysis (LDA), multi-layer perceptron (MLP) and random forest (RF) (Methods). The target (dependent) variable for a chemical was designated as 1 (‘high risk’) if the M/P ratio was ≥ 1, else it was designated as 0 (‘low risk’). Before feature selection, the features with zero variance in the train data were omitted, and thereafter, the dataset was standardized using StandardScaler in scikit-learn (Methods). Subsequently, we performed feature selection using BorutaPy (Daniel, 2022; Kursa and Rudnicki, 2010) to build the classification models (Methods). To optimize our classification models, we performed hyperparameter tuning using GridSearchCV in scikit-learn (Pedregosa et al., 2011) (Methods). For each of the five different classification algorithms, we selected the top 1, 2, 3, 4 and 5 ranked features, and thereafter, trained and evaluated the corresponding model. The best outcome for the five different classification algorithms is reported in Table 1. Notably, the five different classification algorithms were also evaluated using an external test dataset of 202 chemicals which have been experimentally detected in human milk (Methods; Supplementary Table S2; Table 1).

**Table 1:**
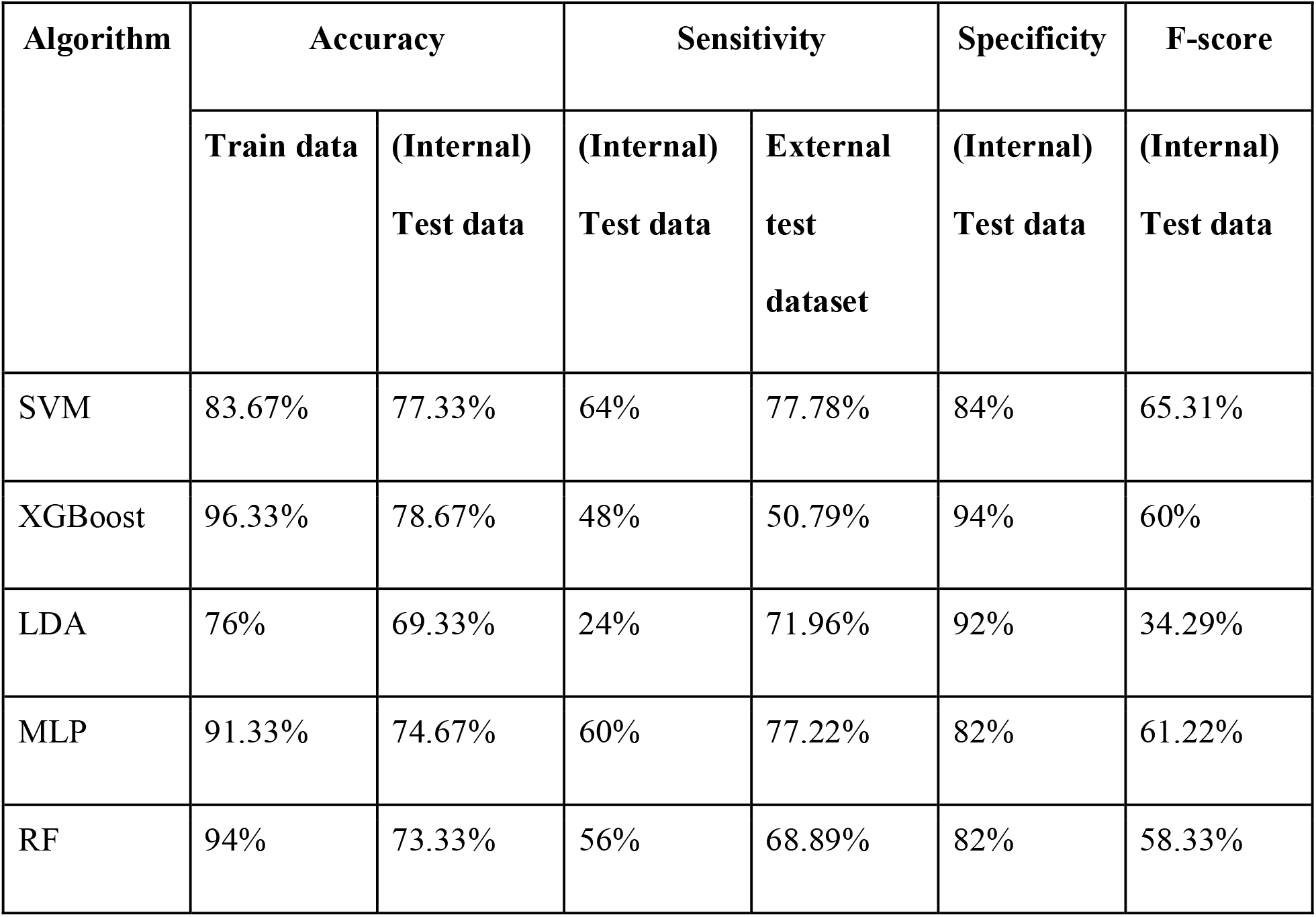
Evaluation of the best classification models built using five different algorithms to categorize chemicals into ‘high risk’ or ‘low risk’ classes for transfer from maternal plasma to human milk. For each model, the classification accuracy is listed for the train data and (internal) test data. The sensitivity is listed for both (internal) test data and the external test dataset respectively. The specificity and F-score are listed for (internal) test data.

A regression algorithm is a statistical method that predicts the continuous values of the dependent variable based on the independent variables (features). To build the regression models to predict the M/P ratio of a chemical, we employed three different machine learning algorithms namely, SVM, XGBoost and RF (Methods). The target (dependent) variable, M/P ratio for a chemical, was logarithmically transformed after adding a constant value of 1 (Methods). Before feature selection, the features with zero variance in the train data were omitted, and thereafter, the dataset was standardized using StandardScaler in scikit-learn similar to classification models (Methods). Subsequently, we performed feature selection using RF (Breiman, 2001) to build the regression models (Methods). To optimize our regression models, we also performed hyperparameter tuning using GridSearchCV in scikit-learn (Pedregosa et al., 2011) (Methods). For each of the three different regression algorithms, we selected the top 5, 10, 15, 20, 25, 30, 35 and 40 features, and thereafter, trained and evaluated the corresponding model. The best outcome for the three different regression algorithms is reported in Table 2.

**Table 2:** Evaluation of the best regression models built using the three different algorithms to predict the M/P ratio of chemicals. For each regression model, we report the coefficient of determination (R^2^) and the Mean Squared Error (MSE) value for both the train data and (internal) test data.

| Algorithm | $R^2$ | | MSE | |
| --- | --- | --- | --- | --- |
|  | Train data | (Internal) Test data | Train data | (Internal) Test data |
| SVM | 0.6909 | 0.4460 | 0.0984 | 0.1978 |
| XGBoost | 0.9323 | 0.4785 | 0.02154 | 0.1862 |
| RF | 0.9277 | 0.4901 | 0.0230 | 0.1820 |

### 3.2 Performance of the classification models

We next evaluated the performance of the five different classification algorithms namely, SVM, XGBoost, LDA, MLP and RF, which were used to build classification models in this study (Methods; Table 1).

In Table 1, we present the accuracy of the classification models on the train data of 300 chemicals and (internal) test data of 75 chemicals. On train data, the models exhibited varying levels of accuracy. Specifically, SVM, XGBoost, LDA, MLP and RF achieved accuracy of 83.67%, 96.33%, 76%, 91.33% and 94%, respectively on train data (Table 1). Moving on to (internal) test data (which is independent of the train data), the five models SVM, XGBoost, LDA, MLP and RF achieved accuracy of 77.33%, 78.67%, 69.33%, 74.67% and 73.33%, respectively (Table 1). Although the accuracy values are relatively high for the five models on (internal) test data, it is important to consider other performance metrics such as sensitivity, specificity and F-score.

Sensitivity measures a model’s ability to accurately predict ‘high risk’ chemicals. On (internal) test data, SVM achieved a sensitivity of 64%, followed by MLP with 60%, RF with 56%, XGBoost with 48%, and LDA with 24% (Table 1). Similarly, specificity measures a model’s ability to accurately predict ‘low risk’ chemicals. On (internal) test data, XGBoost achieved the highest specificity of 94%, followed by LDA with 92%, SVM with 84%, MLP with 82%, and RF with 82% (Table 1). F-score is a measure which combines precision and recall (sensitivity), and thus, provides an overall measure of a model’s performance. On (internal) test data, SVM achieved the highest F-score of 65.31%, followed by MLP with 61.22%. XGBoost with 60%, RF with 58.33%, and LDA with 34.29% (Table 1).

To summarize, in terms of accuracy, sensitivity, specificity and F-score, SVM and MLP showed higher performance on (internal) test data. In contrast, LDA showed lower performance in terms of most evaluation metrics. In comparison, XGBoost and RF achieved reasonable performance but slightly lower than SVM and MLP.

Figure 2 displays the confusion matrix for the five best models corresponding to the five different classification algorithms. The confusion matrix provides the True positive, True negative, False positive and False negative predictions by the five different classification algorithms namely, SVM, XGBoost, LDA, MLP and RF on (internal) test data of 75 chemicals. Thus, the confusion matrix provides a fine print of the performance of the classification models on (internal) test data which concurs with the results presented in Table 1.

**Figure 2:**
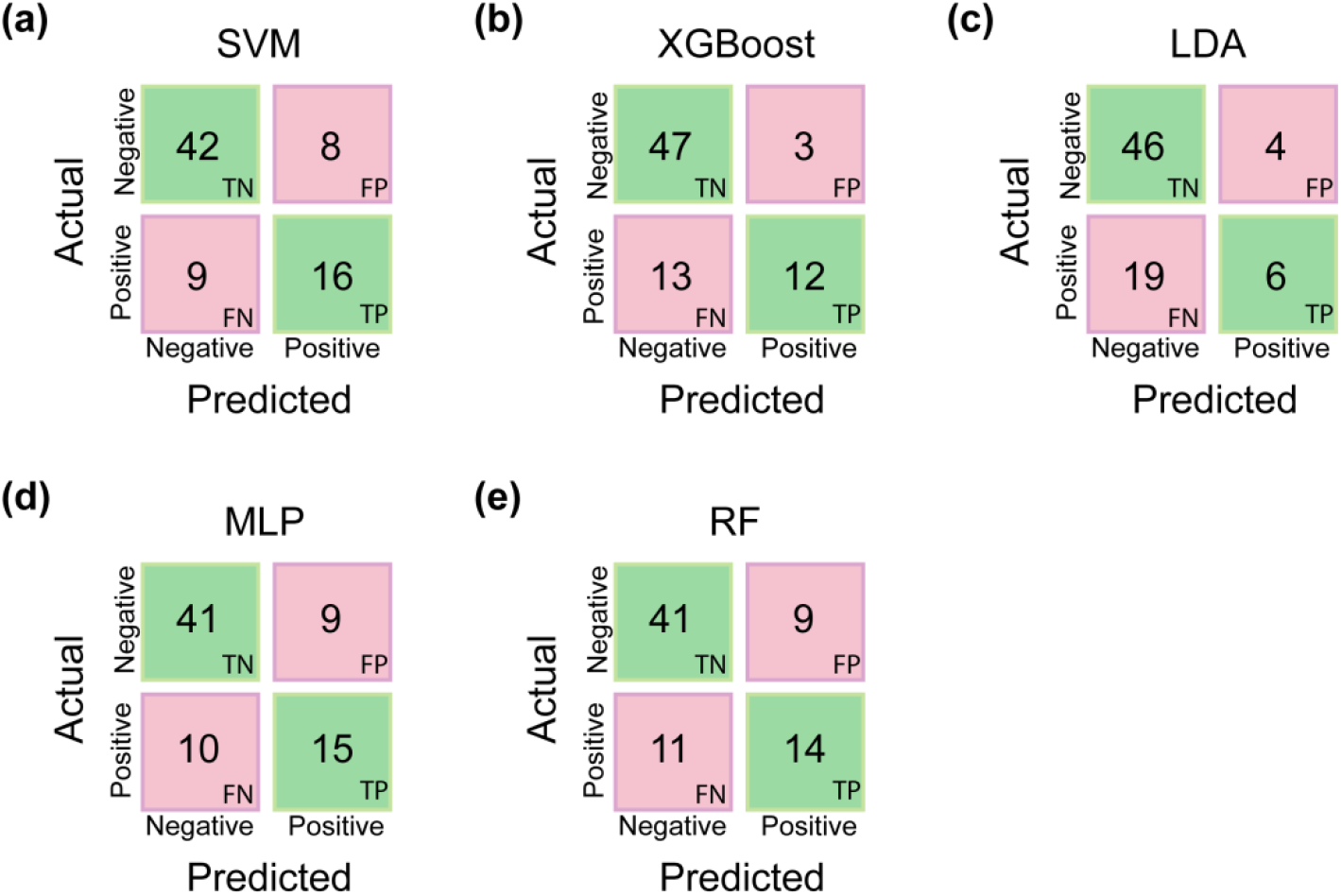
Confusion matrix depicting the performance of the best classification models corresponding to five classification algorithms namely, (a) SVM, (b) XGBoost, (c) LDA, (d) MLP and (e) RF on the (internal) test data of 75 chemicals. Note, TP, TN, FP and FN stand for True Positive, True Negative, False Positive and False Negative.

Furthermore, in addition to the above-mentioned evaluation metrics, we also computed the area under the curve (AUC) values for the five different classification algorithms. Note that the AUC values provide a measure of a classifier’s ability to distinguish between the positive and negative classes in a dataset. Specifically, for the best classification models obtained using SVM, XGBoost, LDA, MLP and RF, the AUC values were 0.82, 0.84, 0.71, 0.77 and 0.84, respectively. These results indicate that XGBoost, RF and SVM classifiers achieve high AUC values, and thus, are better at discriminating between ‘high risk’ and ‘low risk’ chemicals in test data. Figure 3a displays the receiver operating characteristic (ROC) curves for the best models built using the five different classification algorithms.

**Figure 3:**
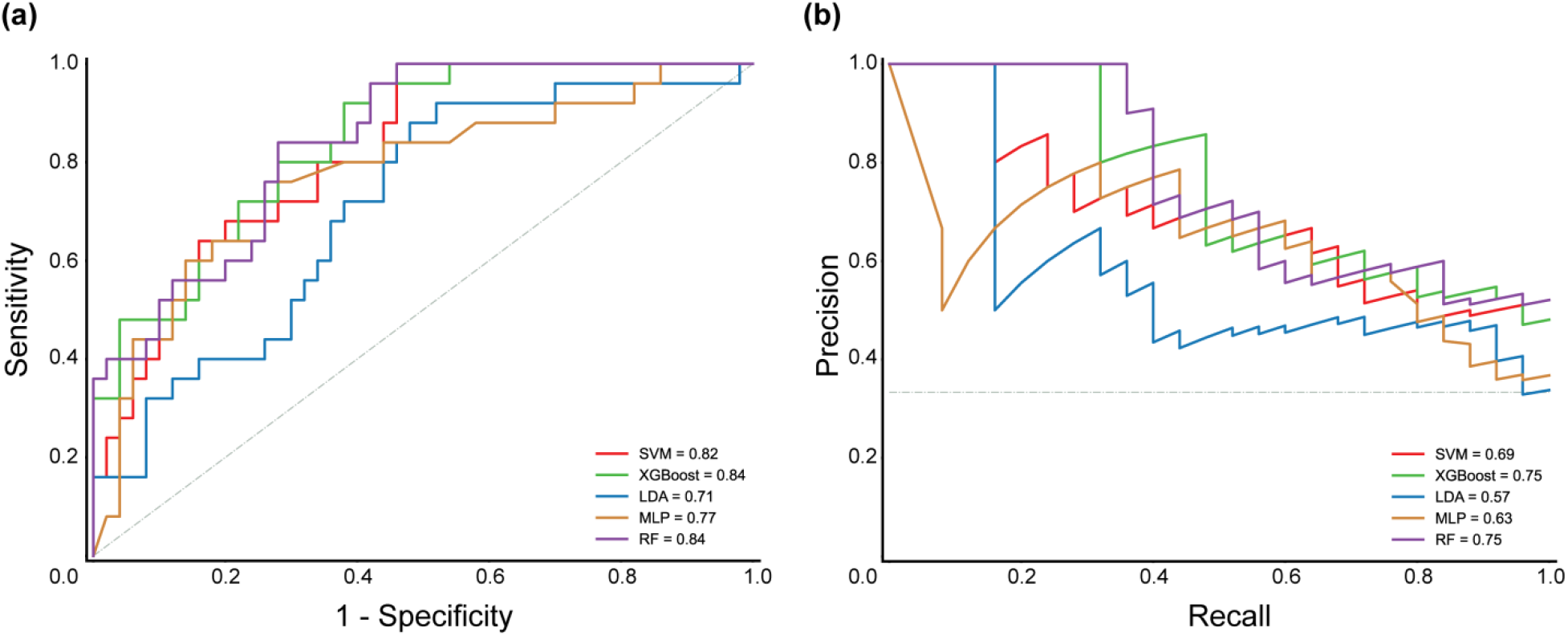
Receiver operating characteristic (ROC) curve and precision-recall curve for evaluating the best models built using the five different classification algorithms. (a) The ROC curve is plotted with sensitivity on the y-axis and 1-specificity on the x-axis. (b) The precision-recall curve is plotted with precision on the y-axis and recall (sensitivity) on the x-axis.

Moreover, we employed the area under the precision-recall curve (AUPRC) as another evaluation metric to assess the performance of the five different classification algorithms. Specifically, for the best classification models obtained using SVM, XGBoost, LDA, MLP and RF, the AUPRC values were 0.69, 0.75, 0.57, 0.63 and 0.75, respectively. These results also indicate that XGBoost and RF classifiers followed by SVM achieve high AUPRC values, similar to the AUC value. Figure 3b displays the precision-recall curves for the best models built using the five different classification algorithms.

Subsequently, we evaluated the generalizability of our best classification models by leveraging an external test dataset, comprising 202 chemicals, with high risk of transfer from maternal plasma to human milk (Methods; Supplementary Table S2). Note that we checked the domain of applicability of data points in the external test dataset before evaluating our classification models (Methods). In Table 1, we present the sensitivity of the five different classification algorithms on external test dataset. By comparing the performance of the five different algorithms on external dataset, it is observed that SVM and MLP achieved the highest sensitivity values of 77.78% and 77.22%, respectively, followed by LDA with 71.96%, RF with 68.89%, and XGBoost with 50.79% (Table 1). In other words, these results indicate that SVM and MLP outperformed the remaining three algorithms in accurately classifying positive instances in the external test dataset. Moreover, the high sensitivities achieved by SVM and MLP on external test dataset suggest that these two algorithms are better capable of capturing the relevant patterns and characteristics associated with positive instances (‘high risk’ chemicals), even though the train data is biased towards negative instances (‘low risk’ chemicals).

Overall, we find that SVM and MLP based classifiers have high predictive ability on both (internal) test data and external test dataset, and thus, the two models are robust and generalizable for the task of accurately predicting high risk chemicals in real-world scenarios (Table 1). Further, among the two models SVM and MLP, we believe that the SVM based classifier is the more promising model for the classification of xenobiotic chemicals into ‘high risk’ or ‘low risk’ of transfer from maternal plasma to human milk.

Importantly, the above-mentioned results also highlight the importance of assessing classifiers on external test datasets to ensure the generalizability of the built models (Bleeker et al., 2003; Cabitza et al., 2021). In particular, such an approach has practical implications while developing reliable machine learning models for drug safety assessment, environmental monitoring, and public health (Bleeker et al., 2003; Cabitza et al., 2021).

### 3.3 Performance of the regression models

We next evaluated the performance of the three different regression algorithms namely, SVM, XGBoost and RF, which were used to build models for the prediction of M/P ratio of xenobiotic chemicals (Methods; Table 2). For the best regression models obtained using SVM, XGBoost and RF, the coefficient of determination (R^2^) values obtained on the train and (internal) test data were (0.6909, 0.4460), (0.9323, 0.4785), (0.9277, 0.4901), respectively. The obtained R^2^ values suggest a moderate level of prediction performance, but also indicate the possibility of overfitting, whereby the models may be fitting the train data more closely than the test data.

To the best of our knowledge, previous studies (Abraham et al., 2009; Agatonovic-Kustrin et al., 2000; Kar and Roy, 2013; Katritzky et al., 2005; Wanat et al., 2020; Yap and Chen, 2005) on building regression-based models for predicting M/P ratio of xenobiotic chemicals have employed much smaller datasets in comparison to our dataset of 375 chemicals with known M/P ratios, and moreover, in many of these published studies (Abraham et al., 2009; Agatonovic-Kustrin et al., 2000; Kar and Roy, 2013), the authors have not reported the R^2^ value on test data. Moreover, none of the previous studies on this topic have published the source code of the developed predictive models. Due to these reasons, it is difficult to compare the performance of our regression models with those reported in the previous studies.

In spite of our attempts to build a highly predictive regression model for the M/P ratio of xenobiotic chemicals, we were unable to improve the R^2^ values on the test data beyond those reported in Table 2 using different regression algorithms. In fact, other than SVM, XGBoost and RF algorithms, we also tried another machine learning algorithm, MLP, to build regression models for prediction of M/P ratio of xenobiotic chemicals. However, MLP yielded highly unsatisfactory regression-based models, and therefore, we decided not to report the results from MLP based regression model in this manuscript. We remark that the difficulty encountered by us in developing a highly predictive regression model for M/P ratio of xenobiotic chemicals has also been faced by authors of previous studies (Abraham et al., 2009; Katritzky et al., 2005), in particular, this issue is well documented in the published literature (Abraham et al., 2009).

While our regression models achieve moderate R^2^ values on (internal) test data, our classification models reported in the previous section achieve high accuracy on (internal) test data. Therefore, we decided to evaluate our best regression models for their ability to perform classification on (internal) test data. Note that if a regression model predicts the M/P ratio of a chemical to be ≥ *log* 2 then the chemical belongs to the ‘high risk’ category, else the chemical belongs to the ‘low risk’ category. We find that our best regression models for SVM, XGBoost and RF achieve a classification based on regression accuracy of 90.67%, 95.33% and 95.33%, respectively on train data, and 74.67%, 72%, and 73.33%, respectively on (internal) test data (Table 3). We also evaluated the sensitivity, specificity and F-score of the classification based on regression models (Table 3). The sensitivity, specificity and F-score for SVM were 60%, 82%, and 61.22%, respectively, for XGBoost were 64%, 76%, and 60.38%, respectively, and for RF were 68%, 76%, and 62.96%, respectively (Table 3). Overall, the results from classification based on regression signify that the models perform reasonably well in categorizing the M/P ratio of xenobiotic chemicals into two classes ‘high risk’ and ‘low risk’ on (internal) test data.

**Table 3:** Evaluation of the classification based on regression models built using the three different algorithms to predict the M/P ratio of chemicals. For each model, the classification accuracy is listed for the train data, the (internal) test data and the external test dataset. The sensitivity is listed for both (internal) test data and the external test dataset. The specificity and F-score are listed for (internal) test data.

| Algorithm | Accuracy |  | Sensitivity |  | Specificity | F-score |
| --- | --- | --- | --- | --- | --- | --- |
|  | Train data | (Internal)<br>Test data | (Internal)<br>Test data | External<br>test<br>dataset | (Internal)<br>Test data | (Internal)<br>Test data |
| SVM | 90.67% | 74.67% | 60% | 60.58% | 82% | 61.22% |
| XGBoost | 95.33% | 72% | 64% | 65.89% | 76% | 60.38% |
| RF | 95.33% | 73.33% | 68% | 75.97% | 76% | 62.96% |

We also evaluated the classification based on regression on external test dataset. Prior to this evaluation, we applied a domain of applicability approach to ensure that external test dataset falls within the range of applicability for our models (Methods). On external test dataset, SVM achieved a sensitivity of 60.58%, XGBoost of 65.89%, and RF of 75.97% (Table 3). These results underscore that the RF model exhibits the highest sensitivity on both (internal) test data and external test dataset, indicating its effectiveness in correctly detecting positive instances (‘high risk’ chemicals). In comparison, the XGBoost model displayed reasonable sensitivity, while the SVM model showed a comparatively lower sensitivity (Table 3).

Overall, we find that the regression models achieve moderate R^2^ values, while the classification based on regression models display reasonable performance on classification of xenobiotic chemicals into ‘high risk’ and ‘low risk’ categories. These results also highlight the complexity and challenges associated with accurate prediction of the M/P ratio of xenobiotic chemicals, and emphasize the need for further research towards creation of improved experimental datasets in order to enhance model performance. Lastly, we remark that none of the three regression based models for classifying the M/P ratio of xenobiotic chemicals, in terms of classification accuracy on (internal) test data and external test dataset, could outperform the SVM based classification model which was solely developed for the classification task (Table 1; Table 3).

### 3.4 Comparison with earlier models

In this subsection, we compare the performance of models built in this study with previously published models for prediction of chemicals with high propensity to transfer from maternal plasma to human milk. Earlier works have employed both classification and regression to make such predictions.

Fatemi and Ghorbanzad’e (2010) (Fatemi and Ghorbanzad’e, 2010) had built a counter propagation artificial neural network based classification model using a train dataset of 124 chemicals. When evaluated on a test dataset of 20 chemicals, this model yielded 100% test classification accuracy (Supplementary Table S5). While this may seem promising, the reported 100% test classification accuracy on a small test dataset necessitates further investigation on the generalizability of this model. Kar and Roy (2013) (Kar and Roy, 2013) had built a LDA based classification model using a train dataset of 97 chemicals. When evaluated on a test dataset of 88 chemicals, this model had test classification accuracy of 56.82%, test sensitivity of 63.16%, and test F-score of 55.81% (Supplementary Table S5). In the present study, we recognized the importance of a larger dataset in improving classification performance. Therefore, we compiled and curated a larger dataset of 375 chemicals with known M/P ratios, of which, 300 chemicals are used as train data and 75 chemicals are used as (internal) test data. Notably, by evaluating our classification models on the (internal) test data of 75 chemicals (which are not part of the train data), we showed that our SVM based classification model achieved test classification accuracy of 77.33%, test sensitivity of 64%, and test F-score of 65.32%, which is a significant improvement over the model developed earlier by Kar and Roy (2013) (Kar and Roy, 2013).

Moving on to earlier publications on regression based prediction of xenobiotic chemicals with high propensity to transfer from maternal plasma to human milk, we find that there is limited reporting of regression R^2^ in such previous studies (Supplementary Table S6). Yap and Chen (2005) (Yap and Chen, 2005) built their model using train data of 102 chemicals and test data of 20 chemicals, and reported a test R^2^of 0.677 and test MSE of 0.206 (Supplementary Table S6). Katritzky *et al*. (2005) (Katritzky et al., 2005) built a QSAR model by dividing a dataset of 100 chemicals into three subsets. Three training datasets were prepared by considering the combinations of any two subsets, and thereafter, equations were obtained for each training dataset. These equations were then used to predict the *log* (M/P) values for the corresponding test datasets. Katritzky *et al*. (2005) (Katritzky et al., 2005) calculated the R^2^ obtained in the three test datasets and reported an average R^2^ of 0.763 (Supplementary Table S6). Wanat *et al*. (2020) (Wanat et al., 2020) built their model using train data of 58 chemicals and test data of 25 chemicals, and reported a test R^2^of 0.29 (Supplementary Table S6). We remark that the above-mentioned three previously published studies developed their regression models using relatively smaller datasets (in comparison to our models), and this may restrict the domain of applicability and limit the generalizability of the three earlier published models. Additionally, the earlier studies also acknowledged the difficulty in achieving high test R^2^values due to limited experimental data and the challenges associated with prediction of M/P ratio for chemicals (Abraham et al., 2009; Katritzky et al., 2005). Among the three regression based models developed in this study, our RF based model achieved a test R^2^ of 0.49 and a mean squared error (MSE) of 0.1820 on (internal) test data (Table 2). Thus, our results align with the limitations mentioned in earlier efforts, wherein previous studies had also encountered challenges in achieving high R^2^ values on test data.

It is also worth noting that many previous studies on this subject do not adequately describe the methods (including technical information) used to build predictive models. This lack of transparency, along with limited reporting of train and test datasets, and unavailability of codes for previous models, hindered direct comparison of our models with earlier works. To facilitate future research in this direction, our study provides detailed documentation of the technical aspects and methods employed to build the predictive models, and moreover, we have made the train data, test data, external test dataset and codes for the built models openly accessible via our GitHub repository: https://github.com/asamallab/M-by-P-ratio-Pred

Overall, the present study highlights the significance of chemical dataset size in building machine learning models for both classification and regression tasks. Further, our study shows the importance of an external test dataset in evaluating the generalizability of built models. By utilizing a much larger train and test data of 375 chemicals plus an external test dataset of 202 chemicals, we created classification- and regression-based models with better performance in terms of predicting the propensity of chemicals to transfer from maternal plasma into human milk.

## 4. Discussion and Conclusion

In this study, we built and evaluated multiple machine-learning models with the aim of classifying xenobiotic chemicals into ‘high risk’ and ‘low risk’ classes for potential transfer from maternal plasma to human milk based on milk/plasma concentration (M/P) ratios of chemicals. We find that our SVM based classifier outperforms other classification models in terms of different evaluation metrics (Table 1; Figure 2). In particular, the SVM based classifier on (internal) test data achieved a specificity of 84%, recall (sensitivity) of 64%, and F-score of 65.31%. Further, from the confusion matrix, it can be seen that the SVM based classifier achieves higher true positives and fewer false positives (Figure 2), and this suggests that the model is less likely to misclassify a ‘high risk’ chemical as a ‘low risk’ chemical. Importantly, based on evaluation on an external test dataset, our SVM based classification model is found to be generalizable, which further strengthens the validity of our approach. In brief, these results attest to the potential of our SVM based classification model to serve as a valuable tool for predicting environmental chemicals whose exposure could pose high risk to maternal and infant health. Notably, such computational toxicology models can serve as valuable alternatives to traditional methods for chemical risk assessment, and have potential to significantly accelerate the pace of characterizing the chemical exposome.

In an effort to support open science we have made our complete workflow, train and test datasets, and computer codes for the built models publicly available via a GitHub repository. We believe this will enable other researchers to reproduce our work without reinventing the wheel, and moreover, facilitate future efforts to leverage our results along with new information to build better predictive models.

While we were able to build highly predictive classification models, we encountered difficulties in developing regression models for predicting the M/P ratio of chemicals. Despite our best efforts, our best regression models for predicting the M/P ratio could achieve only moderate R^2^ values on (internal) test data (Table 2). This issue has been acknowledged by previous efforts (Abraham et al., 2009; Katritzky et al., 2005) in this direction and could be attributed to several factors including the lack of sufficient data to build the models and the influence of environmental factors on the outcome. Interestingly, while the best regression models achieved moderate R^2^ values for predicting the M/P ratio of chemicals, the same regression models performed well in classification based on regression (Table 2; Table 3). However, our best model for classification based on regression could not outperform the SVM model built solely for the classification task. Based on our results, future efforts in this direction may consider exploring different approaches to build separate models for the two tasks, regression and classification.

Overall, our study attests the potential of machine learning models in predicting xenobiotic chemicals with high propensity to transfer from maternal plasma to human milk, while also highlighting the challenges associated with developing accurate regression models for predicting the M/P ratio of such chemicals. While our classification models gave promising results, further investigation is needed to improve the accuracy of regression models. Future efforts to build models with improved prediction power could explore the use of additional features such as genetics, food and lifestyle, physicochemical properties, environmental exposure and toxicokinetic information, or incorporating more sophisticated algorithms such as deep learning.

## Author Contributions

**Sudharsan Vijayaraghavan:** Conceptualization, Data Compilation, Data Curation, Formal Analysis, Software, Visualization, Writing; **Akshaya Lakshminarayanan:** Conceptualization, Data Compilation, Data Curation, Formal Analysis, Software, Writing; **Naman Bhargava:** Conceptualization, Data Compilation, Data Curation, Formal Analysis, Software, Writing; **Janani Ravichandran:** Data Compilation, Data Curation, Writing; **R.P. Vivek-Ananth:**

Conceptualization, Supervision, Formal Analysis, Software, Writing; **Areejit Samal:**

Conceptualization, Supervision, Formal Analysis, Writing.

## Acknowledgements

We thank N. Sukumar for discussions and Kishan Kumar for help with figures. Areejit Samal would like to acknowledge support from the Department of Atomic Energy (DAE), Government of India [Apex project to The Institute of Mathematical Sciences (IMSc), Chennai] and the Max Planck Society, Germany [Max Planck Partner Group in Mathematical Biology]. The funders have no role in the study design, data collection, data analysis, manuscript preparation, or decision to publish.

## Data Availability

The train set, test set and external test dataset along with source codes of the classification- and regression-based machine learning models developed in this study are available via our GitHub repository: https://github.com/asamallab/M-by-P-ratio-Pred

## Supplementary Tables

**Table S1:** Compiled dataset of 375 xenobiotic chemicals with experimentally determined Milk / Plasma concentration (M/P) ratios from published literature. For each chemical, we provide the chemical name, PubChem identifier, CAS identifier, M/P ratio and References in the published literature.

**Table S2:** Compiled list of 202 chemicals in the external test dataset which have been experimentally detected in human breast milk samples. For each chemical, we provide the chemical name, PubChem identifier, CAS identifier and References in the published literature.

**Table S3:** List of best sets of parameters obtained from GridSearchCV for different classification and regression algorithms used in this study to predict the propensity of chemicals to transfer from maternal plasma to milk.

**Table S4:** List of top ranked molecular descriptors (features) for the classification and regression algorithms used in this study for the prediction of the propensity of chemicals to transfer from maternal plasma to milk.

**Table S5:** Methodology and performance metrics for previously published classification models to predict chemicals with high propensity to transfer from maternal plasma into human milk.

**Table S6:** Methodology and performance metrics for previously published regression models to predict M/P ratio of chemicals.

